# A proximity-dependent biotinylation map of cytosol-facing organelle outer membrane proteome in living Arabidopsis cells

**DOI:** 10.1101/2023.03.03.531038

**Authors:** Xinyue Bao, Huifang Jia, Xiaoyan Zhang, Yanming Zhao, Xiangyun Li, Ping Lin, Chongyang Ma, Pengcheng Wang, Chun-Peng Song, Xiaohong Zhu

## Abstract

The cytosol-facing outer membrane (OM) of organelles communicates with other cellular compartments to exchange proteins, metabolites and signaling molecules. Cellular surveillance systems also target OM-resident proteins to control organellar homeostasis and ensure cell survival under stress. Using traditional approaches to discover OM proteins and identify their dynamically interacting partners remains challenging. In this study, we developed an OM proximity labeling (OMPL) system using biotin ligase-mediated proximity biotinylation to map the proximity proteome of the OMs of mitochondria, chloroplasts, and peroxisomes in living Arabidopsis (*Arabidopsis thaliana*) cells. We demonstrate the power of this system with the discovery of cytosolic factors and OM receptor candidates involved in local protein translation and translocation, membrane contact sites, and organelle quality control. This system also performed admirably for the rapid isolation of intact mitochondria and peroxisomes. Our data support the notion that TOM20-3 is a candidate for both a mitochondrial and a chloroplast receptor, and that PEX11D is a candidate for a peroxisome receptor for the coupling of protein translation and import. OMPL-generated OM proximity proteomes are valuable sources of candidates for functional validation and suggest directions for further investigation of important questions in cell biology.

## Introduction

The cytosol-facing membranes of chloroplasts, mitochondria and peroxisomes engage in a wide variety of essential processes, including metabolite and ion exchange, local protein translation and translocation, cellular signaling, organellar biogenesis, and control of organelle quality (Anding and Baehrecke, 2017; Jain and Zoncu, 2022; Reumann and Bartel, 2016; Schleiff and Becker, 2011). High-quality proteome maps of outer membrane (OM) resident proteins—in the outer mitochondrial membrane (OMM), peroxisomal membrane (OMP) and chloroplast membrane (OMC)—and OM-proximal cytosolic factors would be valuable for dissecting the regulatory mechanisms underlying the establishment of both cytosolic and organellar protein homeostasis, as well as organellar quality control. Moreover, a cross-comparison of the OM proximity proteome for the three organelles could identify the proteins located at membrane contact sites that govern inter-organellar communication. However, to date, OMM, OMP and OMC proximity proteomes have not been described in plant cells. One possible reason is that conventional approaches for the enrichment of OM proteins suffer from the loss of large fractions of membrane-resident proteins, together with contamination by proteins from other cellular fractions.

The proximity labeling (PL) approach has demonstrated great potential for probing spatiotemporal interactions between macromolecules and subcellular compartments (Antonicka et al., 2020; Gingras et al., 2019; Hung et al., 2017; Ke et al., 2021; Liu et al., 2018; Mair and Bergmann, 2022; Qin et al., 2021; Yang et al., 2021). In the PL approach, a biotin ligase is fused in-frame with a bait protein, which can be any protein of interest: a cytosolic or nuclear protein or a protein anchored to the membrane of a subcellular compartment. In the presence of biotin, the fused biotin ligase covalently adds a biotin tag to proteins in close proximity to the bait protein. These tagged proteins can be pulled down from extracts using streptavidin beads, allowing researchers to enrich for biotin-labeled soluble proteins and/or insoluble membrane proteins that are usually difficult to obtain by conventional approaches due to their lower abundance or dynamic accumulation patterns. The enriched proteins can then be identified and quantified by classical mass spectrometry (MS).

The PL enzyme TurboID, developed by Ting’s group (Branon et al., 2018), shows fast biotinylation at room temperature using a non-toxic biotin substrate, and the TurboID-based PL technique has been established by several groups in plant systems including Arabidopsis (*Arabidopsis thaliana*), tomato (*Solanum lycopersicum*) root cultures, and *Nicotiana benthamiana* (Arora et al., 2020; Mair et al., 2019; Zhang et al., 2019). These studies have demonstrated that TurboID has a greater efficiency in biotinylating proximal proteins than do other PL enzymes, such as BioID or BioID2, in different plant systems (Conlan et al., 2018; Das et al., 2019; Khan et al., 2018; Lin et al., 2017).

In this study, we establish an OMPL system in Arabidopsis plants. We created transgenic Arabidopsis plants in which TurboID or its truncated variant miniTurboID is genetically targeted to the cytosol-facing OMM, OMP or OMC. TurboID or miniTurboID is fused to a bait protein that is anchored to the surface of the OMM, OMP or OMC, such that the covalent biotinylation of OM proteins and OM-proximal proteins within labeling radius occurs *in vivo* in plant cells when incubated with biotin. Because such biotin labeling is performed at room temperature over few hours while plant cells and organelles are little disturbed, the spatial and temporal relationships between OMs and the proteins in close proximity remain unperturbed. Biotinylated OM proximity proteomes extracted from plant cells represent mostly OM proteins and proximal cytosolic factors.

Using this approach, we examined the OM proximity proteome of three organelles in Arabidopsis cells under normal conditions and ultraviolet-B (UV-B) or high light (HL) stress conditions and tested the feasibility of the PL system for the rapid and specific isolation of intact mitochondria and peroxisomes. We compared our OMM, OMP and OMC datasets and mined each OM proximity proteome in an attempt to identify: 1) candidate cytosolic factors that participate in or regulate organellar protein import; 2) candidate proteins that reside at mitochondrion–peroxisome, mitochondrion–chloroplast, or peroxisome–chloroplast contact sites; and 3) candidate regulatory proteins of the organellar quality control system.

## Results

### Generation of transgenic plants targeting TurboID and miniTurboID to the OM of mitochondria, chloroplasts, and peroxisomes

To develop the PL construct, we used the PL enzyme TurboID or its truncated variant miniTurboID and the OM-anchored proteins OMP25 (the tail-anchored OMM protein synaptojanin-2 binding protein SYNJ2BP) for mitochondria, ATP-BINDING CASSETTE D1 (ABCD1, encoded by At4g39850) for peroxisomes, or TRANSLOCON AT THE OUTER MEMBRANE OF CHLOROPLASTS 64-III (TOC64-lll, encoded by At3g17970) for chloroplasts (Supplemental Table S1). To visualize the localization and to create different labeling radii, we designed two types of constructs: 1) the PL enzyme directly linked to the OM-anchored protein, with green fluorescent protein (GFP) added next to the C-terminal end of the PL enzyme, resulting in constructs named T (TurboID) and Tm (miniTurboID); and 2), GFP inserted between the OM-anchored protein and the PL enzyme to introduce a minimal distance between them, resulting in constructs named TR (TurboID Remote) and TRm (miniTurboID Remote) (Supplemental Fig. S1). We speculated that spacing out the PL enzyme and the OM protein might affect the composition and specificity of the OM proximity proteome because of the resulting change in labeling radius. In addition, in an attempt to achieve fast isolation of intact organelles, we added an HA epitope tag at the C or N terminus of the chimeric protein. In total, we developed four groups of OMPL constructs with TurboID or miniTurboID (Supplemental Fig. S1 and Supplemental Table S1).

We individually transformed these four groups of OMPL constructs into Arabidopsis plants and obtained homozygous transgenic lines: MT-Tm and MT-TRm for mitochondria (MT-OMPL); PX-T and PX-TR for peroxisomes (PX-OMPL); and CP-T and CP-TR for chloroplasts (CP-OMPL) (Fig. 1A). We assessed the correct targeting of each chimeric protein to its intended subcellular compartment in all transgenic lines. We observed that GFP signals from MT-Tm and MT-TRm appeared in leaf and root cells as mitochondrion-like structures; moreover, these GFP signals overlapped with MitoTracker dye-stained dots (Fig. 1B). To check whether PX-T and PX-TR are targeted to peroxisomes, we crossed PX-OMPL plants to a transgenic line harboring a transgene encoding a peroxisome marker fused to mCherry (from the binary plasmid CD3-983). We detected red fluorescence signals from the peroxisome marker around green fluorescence signals in root cells of PX-T and PX-TR seedlings (Fig. 1C), indicating that PX-T and PX-TR were correctly targeted to the OMP in PX-OMPL lines. For CP-OMPL lines, we observed GFP signals in the chloroplast of leaf cells from CP-T and CP-TR plants (Fig. 1D). These results demonstrate that the chimeric proteins for OMPL are targeted to their intended organelles in all examined lines.

**Figure 1.**
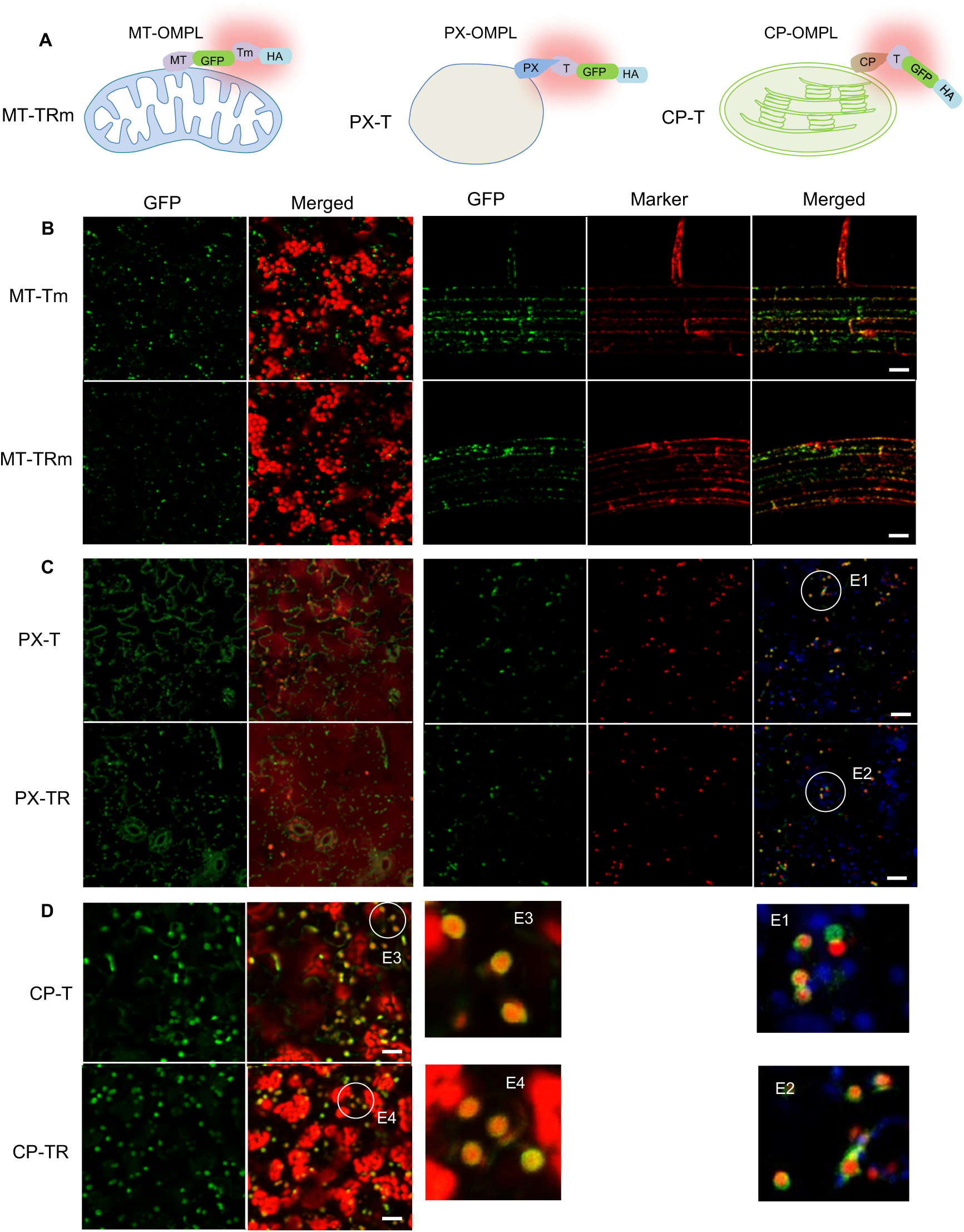
Chimeric proteins containing proximity labeling (PL) enzymes are targeted to specific organellar outer membranes in transgenic Arabidopsis plants. A, Cartoon depicting the designs of organelle outer membrane proximity labeling (OMPL) in this study. MT-OMPL, PX-OMPL and CP-OMPL stand for mitochondrial, peroxisome and chloroplast outer membrane proximity labeling, respectively. T, biotin ligase TurboID; Tm, biotin ligase miniTurboID; MT, PX and CP are mitochondrion, peroxisome and chloroplast outer membrane anchored proteins, respectively; GFP, green fluorescence protein; HA, hemagglutinin epitope. B, Confocal images of transgenic plants expressing *MT-Tm* or *MT-TRm* showing green mitochondrion-like signals in leaf cells (left panel), and co-localization of green mitochondrion-like signals with red mitoTracker signal in root cells stained with MitoTracker™ Red CMXRos. C, Confocal images of transgenic plants expressing *PX-T* or *PX-TR* showing green peroxisome-like signals in leaf cells (left panel), and co-localization of green peroxisome-like signals with red peroxisome marker signal in root cells of *PX-T* or *PX-TR* transgenic plants crossed with peroxisome marker line (CD3-983). Two enlarged images are shown below (E1 and E2). D, Confocal images of transgenic plants expressing *CP-T* or *CP-TR* showing green signals around chloroplasts in leaf cells. Two enlarged images are shown on the left (E3 and E4). See also Supplemental Fig. S1 for the design of OMPL constructs. Scale bars, 20 µM.

To examine the effects of the expression of OMPL constructs on plant growth and response to abiotic stress, we exposed OMPL and wild-type seedlings to various stressors, including antimycin A (AA), an inhibitor of the mitochondrial electron transfer chain that induces mitochondrial stress, and H_2_O_2_, NaCl, and mannitol, which cause oxidative, salt, and osmotic stress, respectively. We checked two or three OMPL lines for each MT-TRm, PX-T and CP-T construct (Supplemental Fig. S2, A–H), and chose one line for each that showed no alterations in response to stress compared to the wild type for analysis.

### Proximity proteomes of OMM, OMP and OMC

To define the proximity proteomes of the three organellar OMs, we optimized conditions for efficient biotin labeling with minimal disturbances of the physiological status of plant cells. We used 14-day-old PX-OMPL seedlings, subjected them to biotin treatment at room temperature, and then monitored biotinylation in total protein extracts using immunoblots with streptavidin-conjugated antibodies. For biotin treatment, we titrated biotin from 0 to 100 μM and tested several durations of treatment. For all tested conditions, we submerged transgenic seedlings alongside the wild-type control in biotin solution at room temperature (approximately 23°C) with shaking. Biotin treatment induced strong labeling of proteins with no major effects on wild-type seedlings, although some proteins were labeled, likely by endogenous biotin ligase activity (Fig. 2A and 2B), revealing the extent of background labeling in this experimental setup. PX-T and PX-TR plants showed different patterns of biotin labeling, suggesting that distance between the PL enzyme and the OM influenced the complement of proteins being labeled by modifying the labeling radius (Fig. 2A and 2B). Given the relatively high labeling efficiency obtained with 14-day-old seedlings treated with 100 μM biotin for 1 h, we used this labeling condition for further analysis. Since TurboID and miniTurboID labeling behaved similarly, as reported previously (Branon *et al*., 2018; Mair *et al*., 2019; Zhang *et al*., 2019), we applied the same labeling conditions to MT-TRm plants in this study.

**Figure 2.**
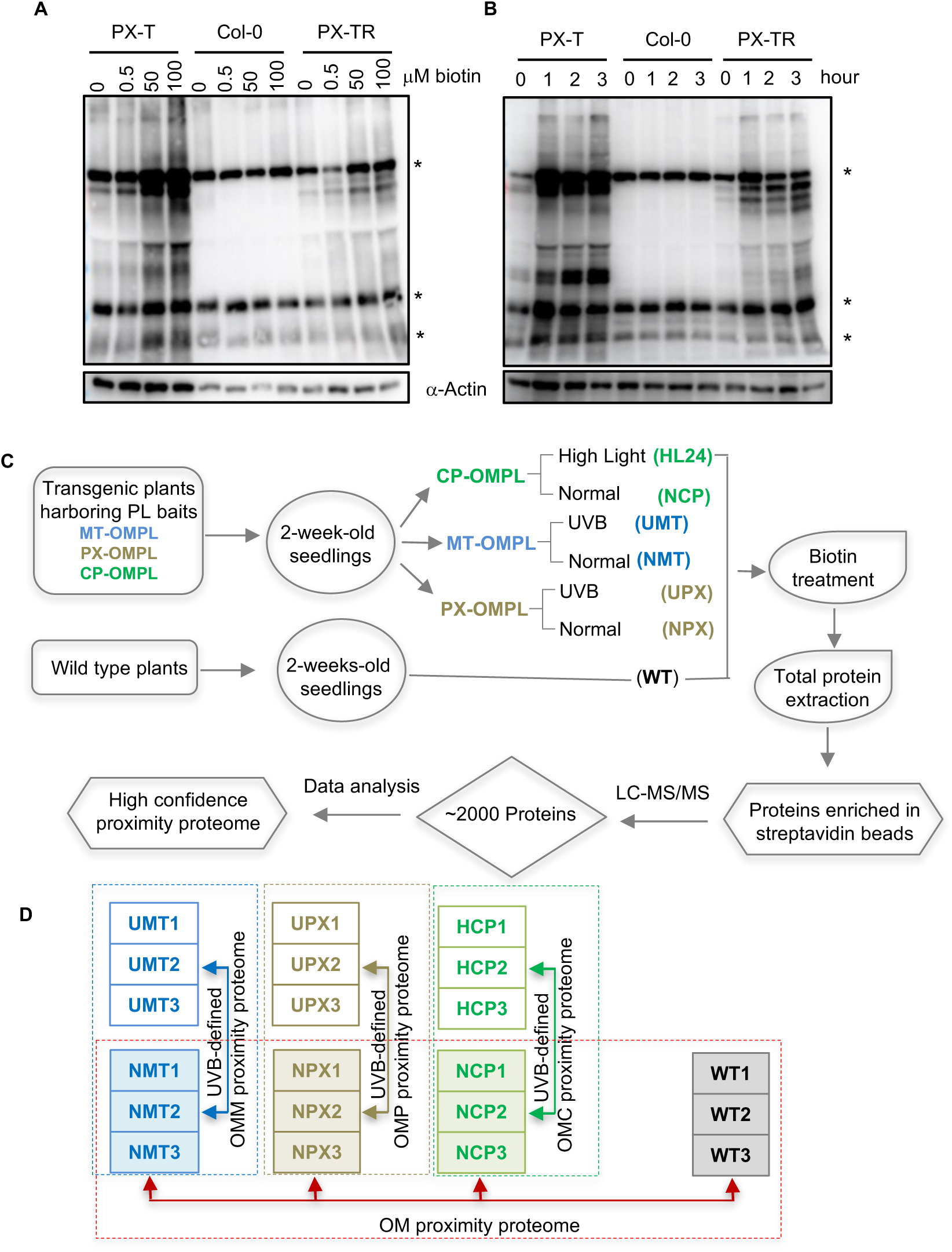
Experimental design for mapping organellar OM proximity proteomes. A and B, Effects of biotin labeling duration and biotin concentration on labeling efficiency. Two-week-old whole Arabidopsis seedlings were submerged in different concentrations of biotin solution as indicated for 1 h (A), or in 100 µM biotin for different durations as indicated (B) at room temperature. Biotin incorporation, reflecting the activity of PL enzymes, was determined by immunoblot with a streptavidin-HRP (SA) antibody, with an anti-actin antibody serving as loading control. Asterisks mark naturally biotinylated proteins. Each sample is a pool of ∼100 mg seedlings. C, Simplified workflow of the experimental procedure from biotin labeling to protein identification by liquid chromatography coupled to mass spectrometry (LC-MS/MS). Three biological replicates were used. For high light and UV-B treatments, two-week-old seedlings were exposed to 2,000 μmol m^-2^ s^-1^ light for 1 h, or 6 w/m^2^ UV for 6 h. High light and UV-B treated seedlings were allowed to recover for 24 h before incubation with biotin. D, Scheme of samples and comparisons used to define OM proximity proteomes under normal growth conditions and following UV-B and HL stress.

We prepared biotin-labeled proteins from OMPL and wild-type (WT) plants grown under normal conditions, and MT-OMPL, PX-OMPL and CP-OMPL protein samples are referred to as NMT, NPX, NCP and WT respectively. All samples were analyzed using liquid chromatography-tandem MS (LC-MS/MS) (Fig. 2C and 2D). We identified 1451 (NMT), 1471 (NPX), 1630 (NCP), and 1706 (WT) high-confidence labeled proteins (defined as being present in at least one sample). Principal component analysis (PCA) showed a separation of the samples by genotype (Fig. 3A). To remove proteins resulting from nonspecific labeling and define the OM proximity proteome, we performed pairwise comparisons between WT and NMT (NMT vs WT), NPX (NPX vs WT), and NCP (NCP vs WT) samples, using only proteins present in all three replicates of the corresponding material. With criteria of absolute fold-change |FC| >1.5 and *p* value <0.05, we retained 311, 273 and 409 OMPL proteins with significantly higher abundance in and specific to NMT, NPX and NCP plants, respectively (Dataset S1). In addition, 57, 42 and 63 OMPL proteins were of significantly lower abundance in NMT, NPX and NCP plants, respectively, compared to WT (Dataset S1).

**Figure 3.**
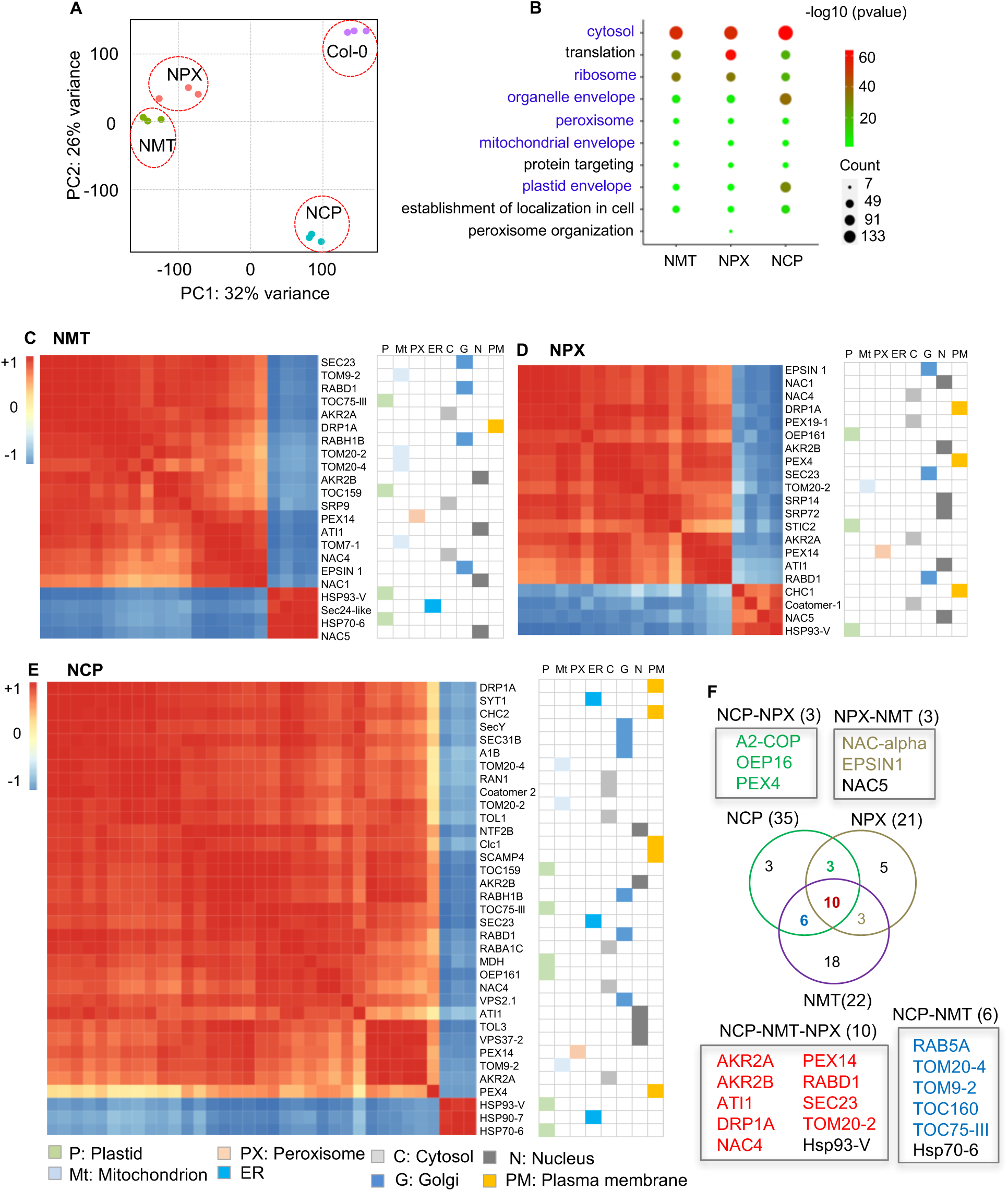
Defining mitochondrial, peroxisome and chloroplast OM proximity proteomes. A, Principal Component Analysis (PCA) showing the clustering of NMT, NPX, NCP and wild-type (WT) samples. B, GO enrichment analysis. Comparison of biological processes and cellular components that are enriched in significantly highly abundant OMPL proteins in NMT, NPX and NCP samples compared to WT samples with filtering criteria of fold-change (FC)>1.5 and *p*-value<0.05. C-E, MT-OMPL (C), PX-OMPL (D) and CP-OMPL (E) Correlation clustering matrices of proteins annotated as “establishment of protein localization”. The subcellular localization of individual proteins based on SUBA5 (the Arabidopsis Subcellular Database) is listed on the right. Colored boxes represent different subcellular compartments. F, Venn diagram showing the extent of overlap between OMPL proteins enriched in the cluster “establishment of protein localization” in NMT, NPX and NCP. OMPL protein names are listed above or below the Venn diagram. Proteins that are not within the same cluster of OM proteins are highlighted in black letters.

To attempt to group the highly abundant proteins from the three OM proximity proteomes in terms of function, we performed a gene ontology (GO) enrichment analysis. The top GO terms in the cellular components category included the ribosome, the cytosol and the organelle envelope; the top GO terms in the biological processes category were related to translation, translation-related processes, and protein targeting (Fig. 3B and Dataset S1). The cellular components ‘mitochondrial envelope’, ‘plastid envelope’ and ‘peroxisome’ were enriched in all three OMPL datasets, while ‘peroxisome organization’ was enriched only in the NPX OMPL set. These results suggest that the three organellar OM baits can capture cytosolic factors in addition to organelle envelope proteins. The three OM proximity proteomes share a common set of subcellular compartment proteins, in addition to organelle-specific proteins.

To identify candidate proteins that might form inter-organellar contact sites, we dissected the GO term ‘establishment of protein localization’ with all significantly altered OMPL proteins in NMT, NPX and NCP samples. This cluster consisted of 22 (NMT), 21 (NPX), and 35 (NCP) proteins, with primary active transporters, small GTPases, chaperones, membrane traffic proteins and subunits of the nascent polypeptide-associated complex (NAC) family of proteins. We predicted the subcellular localization of these proteins based on SUBA5 (https://suba.live), a subcellular localization database for Arabidopsis proteins (Fig. 3C–E). In detail, OMPL detected OM proximity proteins belonging to other cellular compartments in addition to organelle-specific OM proteins. A low correlation of Hsp90-7 and Hsp70-6, Hsp93-V and NAC5 with the list of OM proteins suggested that these cytosolic factors might not be OM proximal proteins. This finding is supported by previous studies demonstrating the cytosolic chaperone Heat shock protein 70 (Hsp70) and Hsp90 have no specific OM docking sites, and act in the post-translational translocation of some soluble proteins (Becker et al., 2019; Schleiff and Becker, 2011). We identified nine proteins common to the NMT, NPX and NCP proximity proteomes: the primary active transporter TOM20-2, PEROXIN14 (PEX14), the chaperones ANKYRIN REPEAT-CONTAINING PROTEIN2A (AKR2A) and AKR2B, the membrane traffic protein DYNAMIN-RELATED PROTEIN 1A (DRP1A), SECRETORY23 (SEC23), the small GTPase RABD1, ATG8-INTERACTING PROTEIN1 (ATI1), and NAC4 (Fig. 3F). Proteins that were present in two of the three OM proximity proteomes included ALPHA2-COP (A2-COP), OUTER ENVELOPE PROTEIN 16 (OEP16) and PEX4 in NCP and NPX; NAC-alpha and EPSIN in NPX and NMT; and RAB5A, TOM20-4, TOM22-V, TOC160 and TOC75-III in NCP and NMT (Fig. 3F). These proteins represent candidates that are involved in inter-organellar contact sites, act as common regulators of organelle protein transport, or have multiple subcellular locations.

To test for possible localization in multiple compartments, we transiently expressed *TOM20-2* and *NAC4* cloned in-frame with the yellow fluorescent protein gene (*YFP*) in *Nicotiana benthamiana* leaves. We determined that the YFP-TOM20-2 fusion appeared to encircle the chloroplast but also formed bright foci, some of which overlapped with the mitochondrial marker or peroxisome marker, suggesting that TOM20-2 may localize to all three organelles (Supplemental Fig. S3, A and B). The localization of TOM20-2 needs to be verified in future studies. By contrast, we observed cytosolic localization of the YFP-NAC4 fusion, as YFP fluorescence did not overlap with either the mitochondrial marker or the peroxisome marker (Supplemental Fig. S3, A and B). Cytosolic NAC4 is thus functionally linked to the OM of three distinct organelles. Taken together, our data demonstrate that the OMPL approach can be used not only to identify mitochondrial, peroxisomal and chloroplast OM proteins but also to catch OM-proximal cytosolic factors. Proteins present in multiple OM proximity proteomes might be candidates for further functional validation, as are proteins specific to each organellar OM proximity proteome.

### UV-B and HL stress affect protein synthesis and import into organelles

To identify proteins that regulate protein import into organelles and are involved in organellar quality control, we subjected MT-OMPL and PX-OMPL plants to UV-B stress (6 W/m^2^ UV for 6 h) and CP-OMPL plants to HL stress (2000 μmol m^−2^ s^−1^ light for 1 h). We then allowed all UV-B-treated MT-OMPL plants (UMT) and PX-OMPL plants (UPX) and HL-treated CP-OMPL plants (HL24) to recover for 24 h before being subjected to biotin labeling and MS analysis, as described above for the NMT, NPX and NCP samples (Fig. 2C and 2D). In total, we identified 1382 (UMT), 1406 (UPX) and 1699 (HL24) high-confidence proteins (in at least one sample). PCA showed a separation between stress-treated and control samples, with a relatively large variance within the group of the three replicates, thus, we used two replicates of NCP and HL24 (Fig. 4A), and two replicates of NMT and NPX, three replicates of UMT and UPX for further comparative analysis (Fig. 4B).

**Figure. 4.**
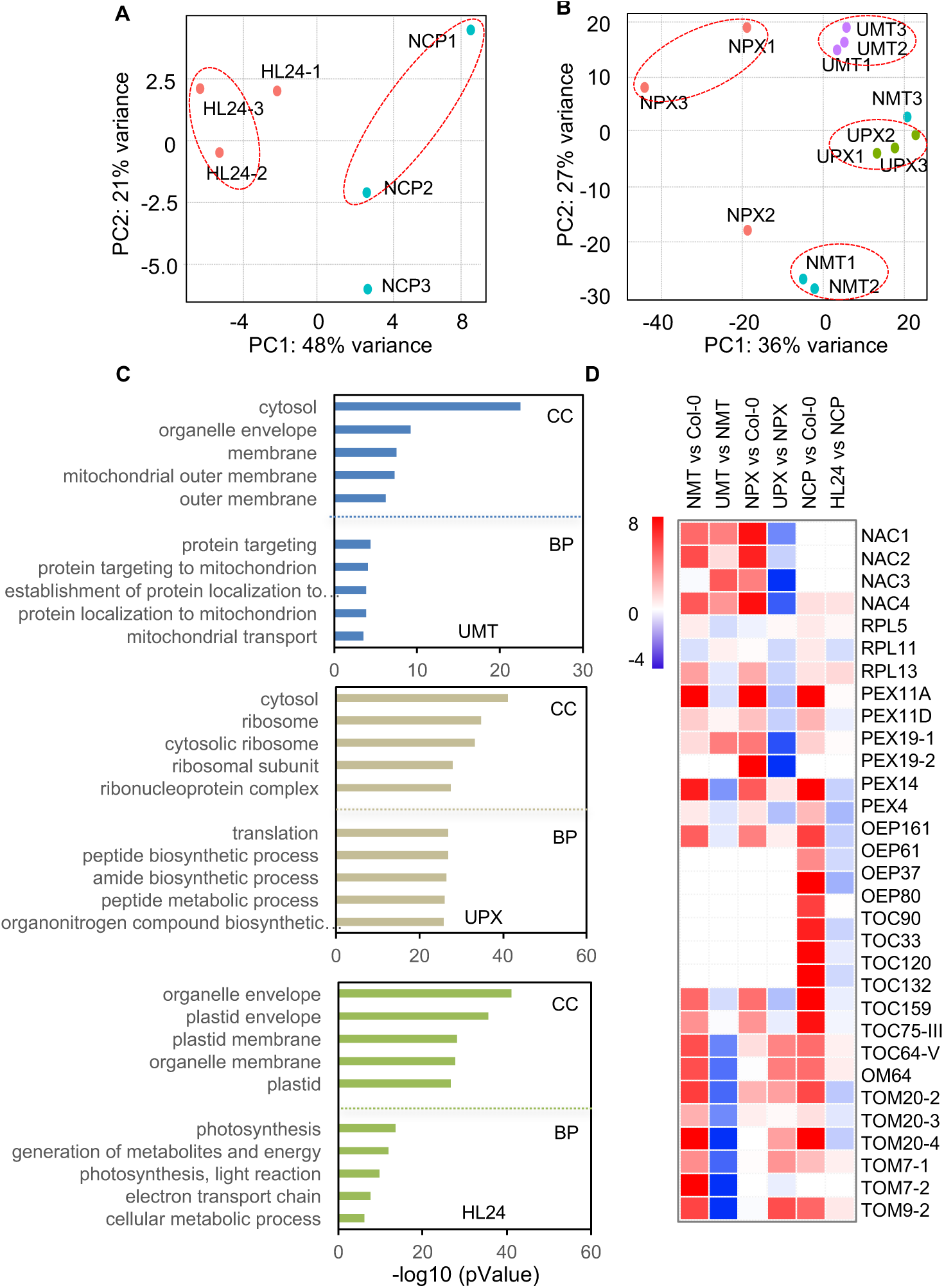
UV-B and HL stress alter the OM proximity proteome. A and B, Principal Component Analysis (PCA) showing the distinction between NCP and HL24 (A), and among NMT and UMT, NPX and UPX samples (B). C, GO enrichment analysis. Top biological processes and cellular component terms enriched with stress-reduced OMPL proteins in UMT, UPX and HL24 samples with filtering criteria of FC<1.5 and *p*-value<0.05. D, Heatmap representation of OMPL proteins with altered abundance in NMT, NPX and NCP compared to WT, and in UMT, UPX and HL24 compared to NMT, NPX and NCP, respectively. Protein names are listed on the right.

To examine whether UV-B or HL stress affects the OM proximity proteome, we performed pairwise comparisons UMT vs NMT, UPX vs NPX, and HL24 vs NCP samples to identify the proteins present in all (two or three) replicates (Fig. 4A and 4B), using the same criteria of |FC| > 1.5 and *p* < 0.05. We retained 88 (UMT), 153 (UPX) and 21 (HL24) proteins whose abundance increased in response to stress and 125 (UMT), 122 (UPX) and 143 (HL24) proteins whose abundance decreased following exposure to stress in OM proximity proteomes (UMT, Dataset S2; UPX, Dataset S3; HL24, Dataset S4). These proteins appeared to be largely specific to UMT, UPX or HL24 samples, as none were detected in all three sets, and only a few (0–10, with a median of 15) were present in two stress-responsive OM proximity proteomes (Supplemental Fig. S4A). These data suggest that UV-B and HL stress each have specific effects on the OM proximity proteome of mitochondria, peroxisomes and chloroplasts. UV-B stress lowered the abundance of proteins related to the biological processes ‘protein targeting’, ‘establishment of protein localization to mitochondria’ and ‘mitochondrial transport’ and the cellular components of ‘organelle envelope’ and ‘mitochondrial outer membrane’ in the UMT samples (Fig. 4C and Dataset S2), suggesting that UV-B stress restrains protein targeting to mitochondria.

We noticed that a large fraction of ribosomal proteins was less abundant following UV-B stress in UPX (Fig. 4C and Supplemental Fig. S4D). In agreement, ‘translation’ and ‘translation-related processes’ were among the top enriched terms among UV-B-repressed proteins in UPX samples (Fig. 4C and Dataset S3). Although ribosomal proteins were not significantly enriched in UMT samples, manual inspection indicated that multiple ribosomal proteins formed two distinct groups, with their abundance either decreasing (Supplemental Fig. S4B) or increasing (Supplemental Fig. S4C) in UMT samples (Dataset S2). These results suggest that UV-B stress influences not only protein synthesis but also protein import into organelles, likely having effects on local protein translation.

Proteins whose abundance declined in response to HL in the chloroplast OM proximity proteome were highly enriched in photosynthesis-related biological processes and organelle envelope and organelle membrane, mostly plastid envelope and plastid membrane (Fig. 4C and Dataset S4), suggesting that HL stress mainly decreases the abundance of photosynthesis-related chloroplast proteins. We manually inspected the proteins in the GO terms ‘protein import’ and ‘establishment of protein localization’ in the OM proximity proteomes obtained after UV-B or HL stress, which revealed that most protein import receptors, for example, TOM20-2, TOM20-3 and TOM20-4, TOM7-1 and TOM7-2, TOM9-2 and OMP64, were significantly less abundant specifically in UMT samples (Fig. 4D). The peroxisome outer envelope pore protein OEP161 and the peroxisome membrane proteins PEX11A and PEX11D were also significantly less abundant in UPX samples (Fig. 4D and Dataset S3). Chloroplast envelope proteins that were specifically enriched in NCP, but not in NMT and NPX samples (for example OEP37, OEP61 and OEP80, TOC90, TOC33, TOC120 and TOC132), appeared to be less abundant, although this result did not reach significance, except for OEP61 and OEP80, which showed a significant decrease in abundance under HL stress (Fig. 4D and Dataset S4). In addition, we observed a significant drop in the abundance of a group of cytosolic chaperone proteins known to function in protein import into organelles (nascent polypeptide-associated complex [NAC] family members NAC1–NAC4, PEX19-1 and PEX19-2, AKR2A and AKR2B) in UPX (Fig. 4D and Dataset S3). NAC1-NAC4, AKR2A and AKR2B were more abundant in UMT (Fig. 4D and Dataset S2). Together, these results suggest that UV-B and HL stress affect the import of nucleus-encoded proteins into organelles, likely having consequences for local protein synthesis in the organelle. The same UV-B stress condition appears to influence chaperone proteins proximal to the OMM and OMP differently.

Few proteins were more abundant upon HL stress, in contrast to the considerable fraction of proteins whose abundance increased in response to UV-B stress in UMT and UPX. UV-B irradiation increased the abundance of proteins located in the chloroplast envelope in UMT, including AKR2A and AKR2B as described above, CLPR3 (ATP-dependent Clp protease proteolytic subunit-related protein 3) and PROBABLE PLASTID-LIPID ASSOCIATED PROTEIN8 (PAP8) (Dataset S2). UV-B stress also caused a rise in the abundance of proteins located at the mitochondrion envelope in UPX, including TOM9-2, VOLTAGE DEPENDENT ANION CHANNEL1 (VDAC1), INORGANIC PYROPHOSPHATASE (TTM2), and NAD KINASE-CAM DEPENDENT (NADKC) (Dataset S3). The abundance of the small GTPases RABA1A, RABE1E, RABH1B, and that of the GTPase activating proteins ARF-GAP DOMAIN6 (AGD6) and AGD9 were increased in UPX, suggesting that UV-B stress likely affects mitochondrion–chloroplast and peroxisome–mitochondrion inter-organellar contacts. Abundance of the autophagy-related proteins ATI1, ATI2, AUTOPHAGY18A (ATG18A), PEX4, PLANT UBX DOMAIN-CONTAINING PROTEIN1 (PUX1), PUX2 and VACUOLAR PROTEIN SORTING34 (VPS34) increased in the peroxisome proximity proteome following UV-B stress, but only ATG18A and PUX1 were significant (Dataset S3). These results suggest that autophagy-related proteins proximal to the OMP were increasingly captured under UV-B stress, although they were not significantly altered in the mitochondrial (Dataset S3) or chloroplast (Dataset S4) OM proximity proteomes under the conditions tested in this study.

### Evidence that TOM20-3 and PEX11D are candidate receptors for local translation coupled with protein import in Arabidopsis cells

We observed a significant enrichment for the cytosolic factors and OM receptors that are involved in protein synthesis and protein transport in the OM proximity proteome, in addition to alterations in the UV-B stress-defined OM proximity proteomes. This result prompted us to examine whether local organellar protein synthesis might be coupled with protein import into organelles in Arabidopsis. We mined the UV-B stress-defined OM proximity proteomes for three types of proteins: import receptor, NAC subunits, and ribosomal proteins. We selected three groups of receptors, TOM20-2 and TOM20-3, PEX11D and PEX11-9 (chaperone as a control), and TOC64 and TOC75III, for mitochondria, peroxisomes, and chloroplasts, respectively. We also chose the NAC subunits NAC1, NAC2, NAC3 and NAC4 and the ribosomal proteins RPS3A, RPL3A, RPL5, RPL11, RPL13, RPL15 and RPL17. Most of these candidates showed a strong enrichment in NMT, NPX and NCP samples, with lower abundance upon UV-B stress or following HL stress (Supplemental Fig. S4, B–D and Fig. 4D).

Using each import receptor, TOM20-2 and TOM20-3, PEX11D and PEX19-1, and TOC75-III and TOC64 as bait, we tested their possible interaction with the NACs and ribosomal proteins by bimolecular fluorescence complementation (BiFC) assay in *N. benthamiana* leaves (Supplemental Table S2). We detected a positive interaction between TOM20-3 and NAC4, RPL5, RPL11 and RPL13, while NAC4 and RPL11 interacted with PEX11D (Supplemental Table S2). The BiFC signals circled chloroplasts when *nYFP-TOM20-3* was co-expressed with *cYFP-NAC4* (Fig. 5A), *cYFP-RPL5* or *cYFP-RPL11* (Fig. 6A) but decorated mitochondria when co-expressed with *cYFP-RPL13* (Fig. 6A). The subcellular BiFC signals were consistent with the dual localization of TOM20-3 in mitochondria and chloroplasts (Fig. 5B). This result is also in line with the OM proximity proteome data showing that TOM20-3 was relatively enriched in NMT and NCP but less abundant in UMT and HL24 (Fig. 4D).

**Figure 5.**
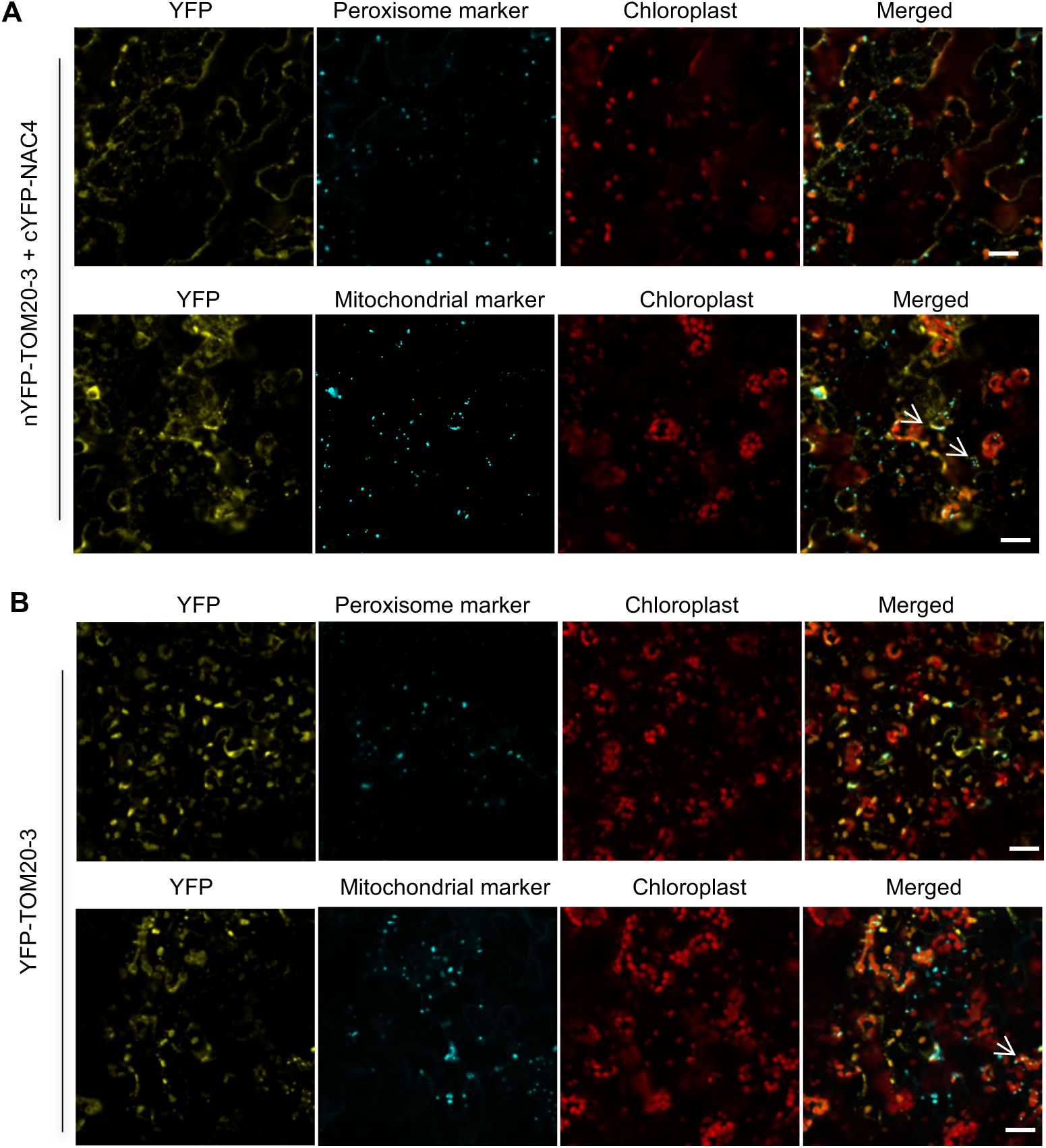
Association of TOM20-3 with NAC4. A, Bimolecular fluorescence complementation (BiFC) assay showing that TOM20-3 interacts with NAC4. BiFC signals surround the chloroplast and appear like mitochondria-like dots, some of which overlap the signal from the mitochondrial marker, but not with the peroxisome marker. B, Confocal images showing that TOM20-3 localizes to both chloroplasts and mitochondria, but not to peroxisomes. Arrows indicate the BiFC or TOM20-3 YFP signals merged with mitochondrial marker CFP signals. Scale bars, 20 μm.

**Figure 6.**
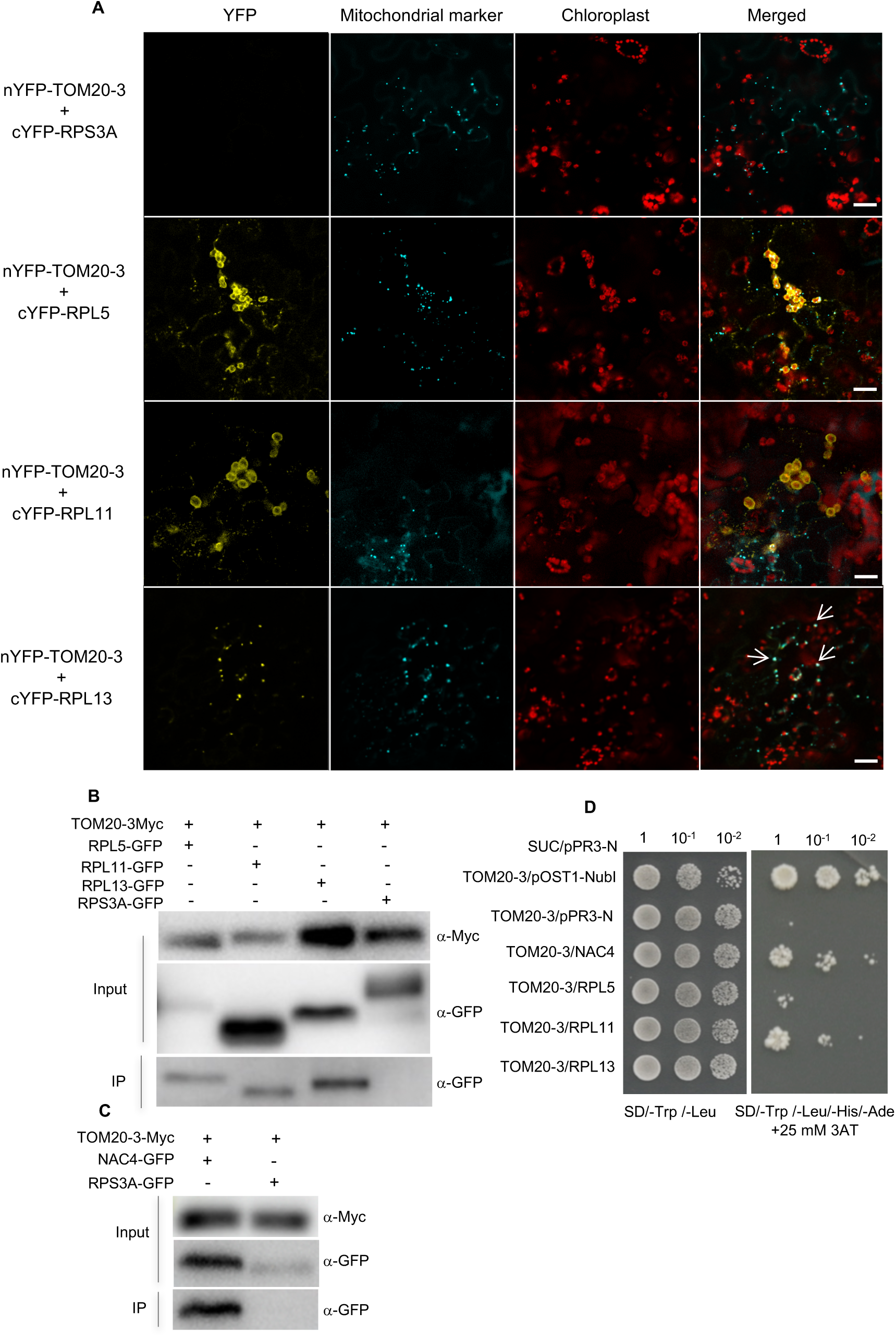
Association of TOM20-3 with ribosomal proteins. A, BiFC assay showing that TOM20-3 interacts with RPL5 and RPL11 around chloroplasts, and with RPL13 around mitochondria. RPS3A was used as a non-interacting negative control. B, Co-immunoprecipitation (Co-IP) assay showing that TOM20-3 interacts with RPL5, RPL11 and RPL13, but not with RPS3A. C, Co-IP assay showing that TOM20-3 interacts with NAC4, but not with RPS3A. D, Yeast two-hybrid (Y2H) assay showing interactions between TOM20-3 and NAC4, TOM20-3 and RPL11, and a relatively weak interaction between TOM20-3 and RPL5. No interaction was detected between TOM20-3 and RPL13. Red fluorescence signals are chloroplast autofluorescence. Arrows indicate BiFC YFP merged with mitochondrial marker CFP signals. Scale bars, 20 μm.

We validated the association of TOM20-3 with NAC4 and the three ribosomal proteins by co-immunoprecipitation (Co-IP) assay (Fig. 6B and 6C). However, a yeast two-hybrid (Y2H) assay failed to detect an interaction between TOM20-3 and RPL13, although TOM20-3 did interact with NAC4 and RPL11 and relatively weakly with RPL5 in this assay, suggesting that the interaction between TOM20-3 and RPL13 may be indirect (Fig. 6D). NAC4 is a cytosolic protein (Supplemental Fig. S3, A and B), but it reconstituted BiFC signals around the chloroplasts and mitochondria with TOM20-3, but not with TOM20-2 (Supplemental Table S2), which also showed a dual localization to mitochondria and chloroplasts (Supplemental Fig. S3, A and B). These results demonstrate that NAC4 interacts specifically with TOM20-3 in the periphery of both mitochondria and chloroplasts.

Following the same procedure, we used PEX11D and PEX19-1 as bait in BiFC assays and established that NAC4 and RPL11 as interacted with PEX11D but not PEX19-1 (Supplemental Table S2). BiFC signals for the PEX11D-NAC4 and PEX11D-RPL11 interactions appeared in peroxisomes, to which PEX11D localized when it was expressed as a *PEX11D-YFP* construct on its own (Fig. 7A). We confirmed the interaction of PEX11D with NAC4, but not with RPL11, by Y2H and Co-IP assays (Fig. 7B and 7C). We also looked for a possible interaction between NAC4 and the three ribosomal proteins by BiFC and Y2H assays. We detected very weak BiFC signals when *cYFP-NAC4* was co-expressed with either *nYFP-RPL5*, *nYFP-RPL11* or *nYFP-RPL13* and restricted to the cytosol (Supplemental Fig. S5). Moreover, we obtained no evidence for a direct interaction between NAC4 and RPL5, RPL11 or RPL13 in a Y2H assay (Fig. 7D).

**Figure 7.**
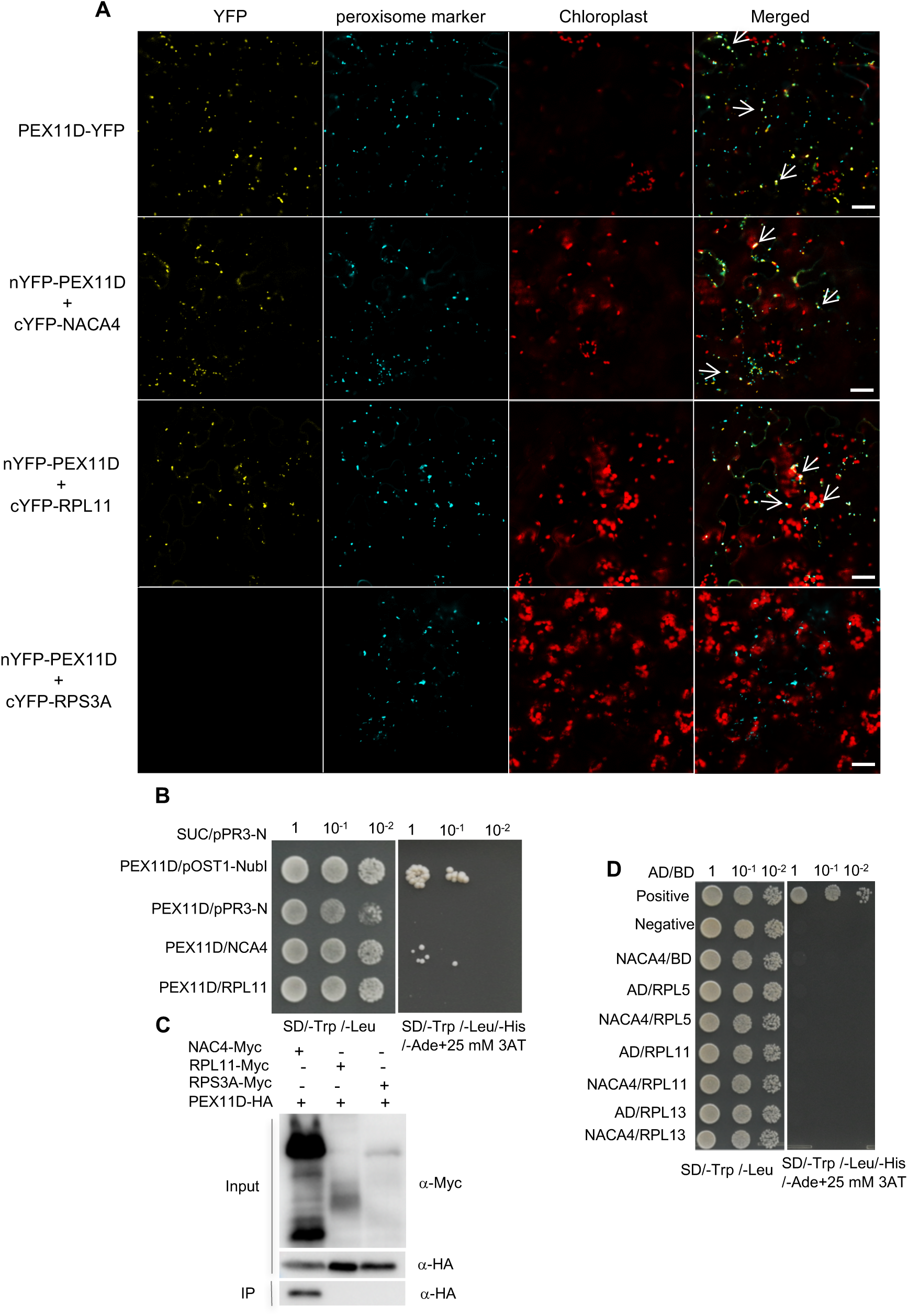
PEX11D interacts with NAC4 and the ribosomal protein RPL11. A, BiFC assay showing that PEX11D interacts with NAC4 and RPL11 around peroxisomes, but not with RPS3A. Arrows indicate BiFC YFP merged with CFP signals from the peroxisome marker. Scale bar, 20 μm. B, Y2H assay showing a relative weak interaction between PEX11D and NPAC4, and no direct interaction between PEX11D and RPL11. C, Co-IP assay showing that PEX11D interacts with NAC4, but not with RPL11 or RPS3A. D, Y2H assay showing that NAC4 does not directly interact with RPL5, RPL11 or RPL13. Scale bars, 20 μm.

Together, these results suggest that TOM20-3 likely acts as both a mitochondrial and chloroplast receptor directly associating with NAC4 and selectively recruits ribosomal proteins. NAC4 may assist the import of newly synthesized proteins into mitochondria, chloroplasts, or peroxisomes.

### Rapid isolation of intact mitochondria and peroxisomes using OMPL plants

The presence of an HA epitope tag at the terminus of chimeric OM proteins enables affinity purification of intact organelles (Boussardon et al., 2020; Chen et al., 2017; Chen et al., 2016; Kuhnert et al., 2020; Niehaus et al., 2020). OMP25 is a well characterized mammalian mitochondrial protein that was previously used for isolating intact mitochondria (Chen *et al*., 2016). We validated the mitochondrial localization of chimeric OMP25 in MT-OMPL plant cells (Fig. 1A). We noticed no mistargeting of the OMP25 chimera to chloroplasts in our study. ABCD1 is a peroxisomal protein and a member of the Arabidopsis ABC transporter family. To generate PX-OMPL transgenic lines, we used the N-terminal first 100 amino acids of ABCD1 to target our chimeric protein to peroxisomes (Fig. 1B). Again, we observed the correct localization of the resulting chimeric protein in OMPL plants.

We followed a similar procedure for the fast isolation of intact mitochondria and peroxisomes, using magnetic affinity resin decorated with anti-HA antibodies as described previously (Chen et al., 2017). To enrich for HA-tagged mitochondria from MT-OMPL plants and HA-tagged peroxisomes from PX-OMPL plants, we collected 1-week-old MT-OMPL and PX-OMPL whole seedlings that had been grown vertically on Murashige and Skoog (MS) plates. Our purification procedure consisted of five steps: 1) grinding of seedlings (1 min); 2) centrifugation of the homogenates (5 min); 3) incubation of the supernatant with HA beads (10 min); 4) collection of HA beads (1 min); 5) washing of beads (3 x 1min) (Fig. 8A). This method allowed us to purify mitochondria and peroxisomes from plant material in about 20 min (Fig. 8A). We inspected the purified mitochondria and peroxisomes for their intactness under a confocal microscope; we also tested the purity of each bead preparation by immunoblot analysis with organelle-specific marker proteins. Only HA beads incubated with homogenates prepared from MT-OMPL and PX-OMPL seedlings showed green fluorescence under the microscope; those incubated with extracts from wild-type nontransgenic seedlings did not. These results demonstrate the specific enrichment of intact mitochondria or peroxisomes (Fig. 8B).

**Figure 8.**
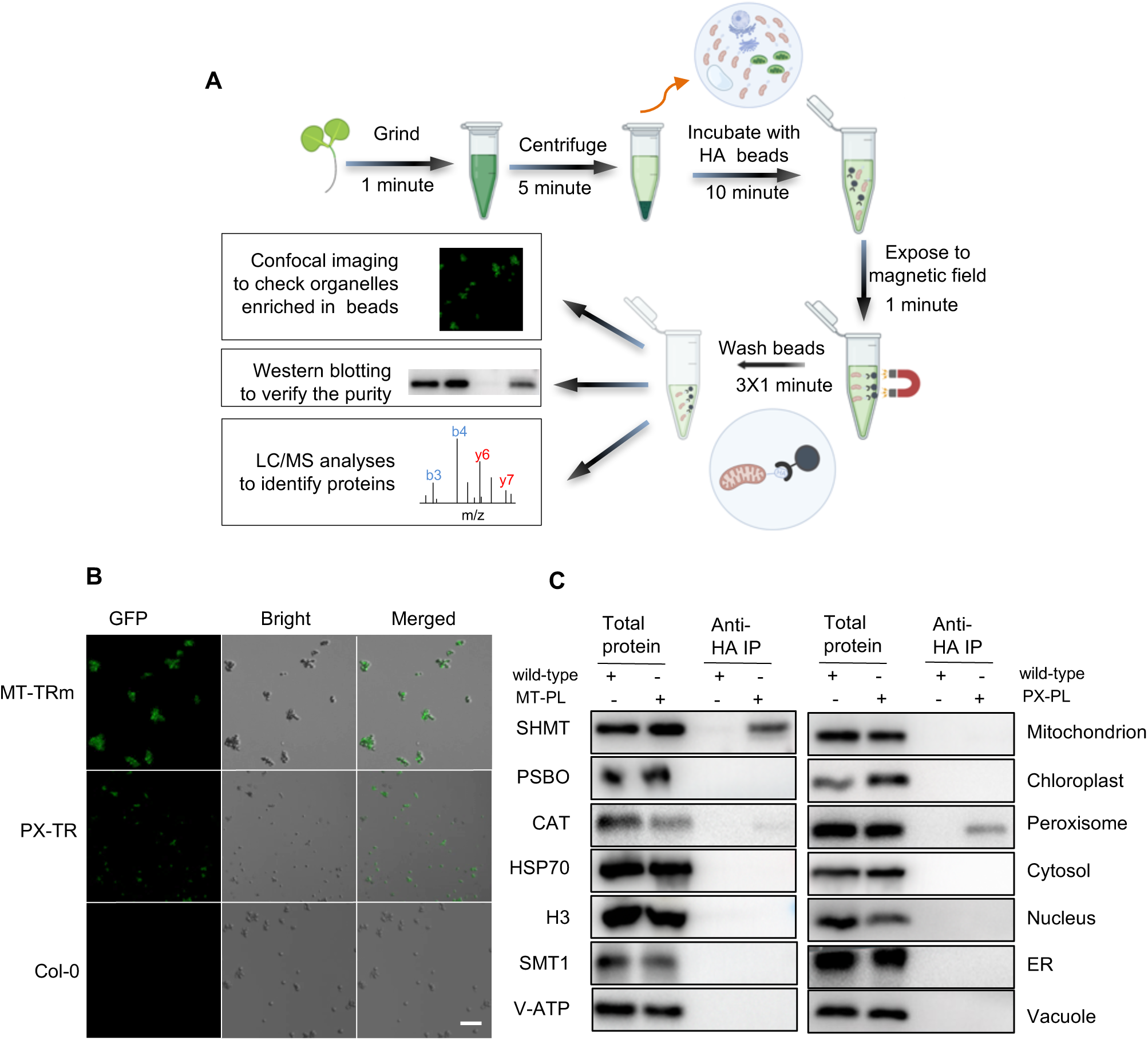
Rapid isolation of intact mitochondria and peroxisomes from OMPL plants. A, Workflow for the isolation of intact mitochondria and peroxisome. B, Confocal images showing green signals from intact mitochondria and peroxisomes bound to HA beads. C, Immunoblot showing potential contaminations with other cellular compartments from isolated intact mitochondria (left panel) and peroxisomes (right panel). Antibodies used for detecting each compartment are indicated next to panels. Scale bar, 5 μm.

Similarly, immunoblot analyses confirmed the enrichment for mitochondria and peroxisomes, respectively, from MT-OMPL and PX-OMPL plants, but not from the wild type, using mitochondrial matrix protein serine hydroxymethyltransferase (SHMT) as a mitochondrial marker and catalase (CAT) as a peroxisomal protein. Using antibodies against PSBO (33 kDa subunit of oxygen evolving system of photosystem II, encoded by At3g50820, plastid marker), HSP70 (cytosol marker), STEROL METHYLTRANSFERASE1 (SMT1, endoplasmic reticulum [ER] marker), histone H3 (H3, nucleus marker), and V-ATP (V-ATPase, vacuolar marker), we observed minimal contamination from other cellular compartments in our mitochondria- and peroxisome-enriched beads, demonstrating the low extent of contamination from these cellular compartments (Fig. 8C). These results suggest either contamination of one organellar preparation by the other or a physical interaction between mitochondria and peroxisomes. We observed no enrichment for mitochondria or peroxisomes on beads incubated with extracts prepared from the wild type, indicating that the beads bind specifically to HA-tagged mitochondria or peroxisomes in the MT-OMPL or PX-OMPL lines, respectively (Fig. 8B and 8C). These results also verified that the HA-TurboID–GFP-OMP25 and PX-TurboID-GFP-HA chimeric proteins were anchored to the surface of mitochondria and peroxisomes, respectively, and point towards the cytosol. Taken together, our data indicate that intact mitochondria and peroxisomes can be rapidly and specifically isolated via the methods established with MT-OMPL and PX-OMPL plants. The OMPL approach combined with this rapid isolation of intact organelles should be a powerful tool for dissecting seminal questions related to organellar protostasis control, metabolite exchange, and inter-organellar communication.

## Discussion

Here, we describe our efforts to map the OM proximity proteomes on the cytosolic sides of mitochondria, peroxisomes and chloroplasts in plant cells via stable transgenic Arabidopsis plants expressing constructs encoding the PL enzyme fused to specific OM-anchored proteins. We compared the abundance of each MS-detected protein in OMPL plants and wild-type plants. We defined three OM proximity proteomes, which are mainly composed of cytosolic factors and organellar envelope proteins involved in protein translation, protein targeting and establishment of protein localization. This similarity suggests some commonalities in biological processes in addition to features specific for each OM proximity proteomes, which included specific responses to UV-B or HL stress. All proteins identified here in the OM proximity proteomes define 1) potentially novel cytosolic factors or OM proteins participating in local protein synthesis and/or import of proteins into mitochondria, peroxisomes and chloroplasts; 2) components forming mitochondrion–chloroplast, mitochondrion–peroxisome, or chloroplast– peroxisome contact sites; 3) regulators of organellar quality; and 4) proteins with multiple subcellular locations. Hence, OM proximity proteomes are valuable resources for advancing our understanding of the regulatory machinery governing these three important organelles and for generating novel hypotheses in plant cell biology.

To demonstrate the application of our OM proximity proteomes in a biological discovery context, we focused on protein transport to organelles, which is functionally linked to organellar and cytosolic protostasis, as well as organellar biogenesis (Gao et al., 2023; Topf et al., 2016; Wrobel et al., 2015). Mitochondria, peroxisomes and chloroplasts contain several hundred proteins that are critical for their functions, most of which are encoded in the nucleus, synthesized in the cytoplasm, and imported post-translationally into each organelle through organellar import complexes present in their envelopes. Although this post-translational import mechanism has been established for mitochondria, peroxisomes and chloroplasts in plant cells, the detailed mechanisms remain elusive (Baker et al., 2016; Cross et al., 2016; Fish et al., 2022; Lashkevich and Dmitriev, 2021; Schleiff and Becker, 2011; Topf et al., 2019).

Proteins can also be imported co-translationally while being synthesized, a concept that was revisited by recent studies of mitochondria and peroxisomes in yeast (*Saccharomyces cerevisiae*) (Becker et al., 2019; Dahan et al., 2022; Gold et al., 2017; Lesnik et al., 2014; Lesnik et al., 2015; Williams et al., 2014) and in human cells (Gehrke et al., 2015). To the best of our knowledge, this co-translational import mechanism has never been reported in plant cells. Even in yeast, several questions remain: 1) What is the relative contribution of the two mechanisms–post-translational import and co-translational import–in organellar protein import? 2) What is the specificity of protein targeting by each mechanism, if any? For example, are membrane proteins preferentially targeted by co-translational import while matrix proteins are targeted by post-translational import? 3) Do these two mechanisms respond differently to developmental and stress cues?

NAC is a ribosome-associated chaperone that interacts with newly synthesized proteins as they exit the ribosome, and its function has been implicated in protein co-translational targeting to organelles (Albanese et al., 2010; del Alamo et al., 2011; Funfschilling and Rospert, 1999; Kirstein-Miles et al., 2013; Kramer et al., 2019). In the mode of NAC action in co-translational import, the association of NAC with organelles necessitates an OM receptor. OM14 is a primary receptor that is critical for the association of NAC with mitochondria during mitochondrial protein co-translational import in yeast (Lesnik et al., 2014). However, a receptor for NAC on the mitochondrial, peroxisomal or chloroplast outer membrane has not been identified in plant cells.

Our study provides three critical pieces of evidence that establish co-translational import in mitochondria, peroxisomes, and chloroplasts in plants. First, we identified candidates for the protein import receptors: TOM20-3 for NAC on the mitochondrial and chloroplast outer membranes and PEX11D for NAC on the peroxisomal outer membrane. The evidence supporting TOM20-3 acting as both a mitochondrial and chloroplast OM receptor includes the following: 1) the dual mitochondrial and chloroplast localization of TOM20-3 (Fig. 5B); 2) the physical association of TOM20-3 with NAC4 in mitochondria and chloroplasts (Fig. 5A); and 3) the direct interaction of TOM20-3 and NAC4 in Y2H (Fig. 6C and 6D). Along the same lines, we propose that PEX11D is a candidate peroxisome receptor that associates with NAC based on the following evidence: 1) the peroxisomal localization of PEX11D (Fig. 7A); 2) the association of PEX11D with NAC4 in peroxisomes (Fig. 7A); and 3) the direct interaction of PEX11D and NAC4 in Y2H (Fig. 7B and 7C). Second, we detected NAC4 in all three OM proximity proteomes, where it likely acts as a multifunctional chaperone and plays a role in co-translational protein import. Its functional plasticity may be achieved by the recognition by its OM receptor and its client proteins. Third, we showed that UV-B and HL stress altered the abundance of these candidate OM receptors, NAC subunits and ribosomal proteins, suggesting that local translation and translocation respond to stress conditions.

Previous studies suggested a function for TOM20-3 in mitochondrial protein import based on the following evidence: 1) TOM23-1 directly interacted with mitochondrial precursor proteins; 2) the overexpression of Arabidopsis *TOM20-3* in yeast resulted in competition for protein import into yeast mitochondria (Lister et al., 2007); 3) a nuclear magnetic resonance structural analysis revealed a very similar presequence binding fold in TOM20-3 and its mammalian ortholog TOM20 (Likic et al., 2005; Perry et al., 2006); and 4) the rate of protein import was lower in a *tom20-2 tom20-3 tom20-4* triple knockout mutant (Lister et al., 2007). Our findings, combined these previous results, suggest that TOM20-3 may be involved in both post-translational import and co-translational import.

Arabidopsis has five PEX11 proteins, PEX11A to PEX11E, that are all targeted to peroxisomes and promote peroxisome proliferation (Orth et al., 2007). We detected an interaction between PEX11D and NAC4 and RPL11 in peroxisomes, although we observed no evidence for a direct interaction for PEX11D–RPL11, which could be a very weak interaction that falls below the detection limit of the method used. PEX11D mediating co-translational targeting raises the possibility of a functional link between co-translational protein import and peroxisome biogenesis in plant cells, which is an interesting question for future study. Co-translational targeting specificity for peroxisomal membrane proteins (PMPs) was shown by a study in yeast (Dahan et al., 2022). However, how localized translation is coupled with translocation of PMPs to peroxisomes has not been established. We demonstrated that a fraction of ribosomal proteins is proximal to peroxisomes and that their abundances were significantly reduced under UV-B stress. The presence of ribosomal proteins on the surface of peroxisomes can be examined by transmission electron microscopy in future studies to look for similarities with mitochondria (Gold et al., 2017). Our findings provide a ground for further investigations.

The discovery of AKR2A and AKR2B in all three OM proximity proteomes suggests that these two Arabidopsis ankyrin repeat proteins may function as cytosolic chaperones not only for sorting and targeting chloroplast OM proteins to the chloroplast, as shown in previous studies (Bae et al., 2008; Kim et al., 2014), but probably also for mitochondrial and the peroxisomal proteins. This finding is in line with previous work demonstrating that AKR2A is a molecular chaperone for a group of proteins that contains an AKR2A-binding sequence (ABS). This group of ABS-containing proteins include the chloroplast outer envelope proteins OEP7 and TOC34, the peroxisomal membrane-bound proteins ASCORBATE PEROXIDASE3 (APX3) and APX5, the mitochondrion outer membrane protein TOM20-4 and the microsomal membrane (ER-membrane) proteins CYTOCHROME B_5_ (B FORM OR CB5-B) and CYTOCHROME B_5_ REDUCTASE (CB5R) (Zhang et al., 2010). Therefore, AKR2A and AKR2B are likely multifunctional chaperones for sorting and targeting membrane proteins to different cellular compartments. Given that AKR2A-mediated targeting of chloroplast OM proteins is coupled to their translation (Kim et al., 2015), the possibility of AKR2A-TOC34-RPL23 mediated local translation awaits future investigation. Moreover, our findings also raise the possibility that mitochondrial and peroxisomal membrane proteins are targeted to their respective organelles by AKR2A or AKR2B, using a mechanism similar to that of AKR2A-mediated protein targeting to chloroplasts (Bae et al., 2008; Kim et al., 2014). If this is the case, the regulation of AKR2A- and/or AKR2B-mediated protein targeting in response to stress would be an interesting focus for future studies, given that the abundance of AKR2A and/or AKR2B proximal to organellar OM is subjected to regulation under stress conditions.

The application of the OMPL system for the rapid and specific isolation of intact mitochondria and peroxisomes that we developed would be useful in combination with OMPL for discovering the biological function of each component, for example, in cell biology of protein targeting, in inter-organellar communication of metabolite exchange, and in organellar quality control during cell senescence and response to various stresses. The rapid and specific isolation of intact organelles would enable spatiotemporal tracking of organelle proteomes, metabolomes and transcriptomes that can largely reflect the situation *in vivo* using minimal sample materials. Thus, OMPL would be a better choice than other approaches, such as density gradient centrifugation, time-consuming and not very specific.

Limitations of our study should be noted here. 1) The current approach was unsuccessful for the specific isolation of intact chloroplasts, probably due to their large size. Although we achieved a relative enrichment of chloroplasts, the specificity and purity were not guaranteed. 2) We often co-purified mitochondria and peroxisomes, which might be the result of mitochondrion–peroxisome physical interactions, as well as possible contamination. Despite this limitation, a high enrichment can still be achieved for mitochondria or peroxisomes. 3) The proximity proteomes defined in this study may exclude important proteins for four reasons. a) The 14-day-old seedlings used in this study may not be suitable for identifying OM proximity proteins that participate in the plant reproductive stage. b) The relatively large variation among replicates introduced during biotin labeling and stress treatments would prevent the identification of important proteins. For example, the significant level of stress-induced autophagy-related proteins often fell below the threshold because of large variations. Thus, consistent conditions for biotin and stress treatments are critical for maximizing informative outcomes. c) A fraction of non-specific proteins may exist in our OM proximity proteomes. d) Although we created constructs with different labeling radii and two versions of the PL enzyme, TurboID and miniTurboID, we were not able to compare the effects of the labeling radius, or compare TurboID and miniTurboID on the composition of the OM proximity proteome, due to experimental cost. Nevertheless, the choice of plant material and application of biotin labeling conditions and stress treatments should be optimized based on the focus of the study.

In conclusion, our study established mitochondrial, peroxisomal and chloroplast OMPL systems in Arabidopsis plants and produced six high-quality OM proximity proteomes for plant mitochondria, peroxisomes and chloroplasts, demonstrating that OM proximity proteomes can be mined for novel protein candidates in organellar co-translation and translocation, organelle membrane contact sites, and organellar quality control.

## MATERIALS AND METHODS

### Plant materials and growth conditions

All wild-type and transgenic Arabidopsis (*Arabidopsis thaliana*) lines used in this study were in the Columbia (Col-0) accession. All Arabidopsis seedlings used for the analyses of the growth phenotypes were grown on half-strength MS (Caisson labs) solid medium with 1% (w/v) sucrose or on MS medium solidified with 0.8% (w/v) agar or 0.6% (w/v) Phytagel and containing stressors when appropriate. Plates were placed in growth chambers at 22°C under 100 μmol m^−2^ s^−1^ light (WT5-LED14, FSL) in a 16-h light/8-h dark photoperiod.

### Generation of constructs and OMPL transgenic plants

Plasmids carrying the sequences encoding the PL enzymes TurboID (T) and miniTurboID (Tm) were gifts from Dr. Dinesh-Kumar SP (UC Davis); the *OMP25* cDNA was purchased from Addgene (https://www.addgene.org). Arabidopsis cDNAs for *ABCD1* (At4g39850) and *TOC64-lll* (At3g17970) and the coding sequences of various genes for BiFC, subcellular localization, Co-IP and Y2H analysis were all amplified by RT-PCR using gene-specific primers (Dataset S3). The fragments of *TurboID* or *miniTurboID* and *OMP25* or *ABCD1* or *TOC64-III* were cloned in-frame into the binary vector backbone 1300-EGFP-3XHA or 1300-Flag or 1300-EGFP (Supplemental Fig. S1 and Supplemental Table S1) using one step cloning kit (ClonExpress II One Step Cloning Kit | #C112, Cellagen Technology). The vector backbones for BiFC constructs were pSITE-nYFP-C1 (CD3-1648), pSITE-cYFP-C1 (CD3-1649), pSITE-nYFP-N1(CD3-1650) and pSITE-cYFP-N1(CD3-1651) from ABRC. The constructs for subcellular localization and Co-IP analysis were generated using the Gateway vectors pDONR207, pEARLEYGATE101 (CD3-683), pEARLEYGATE104 (CD3-686) and pEARLEYGATE203 (CD3-689) from ABRC. The constructs for Y2H analysis were generated for OM proteins using the vectors pBT3-N, pBT3, pBT3-SUC and pBT3-STE from the split ubiquitin yeast two-hybrid system and for cytosolic proteins using pGBKT7:BD and pGADT7:AD from the nuclear yeast-two hybrid system DUAL hunter. Transgenic plants were generated by transforming OMPL constructs (Supplemental Fig. S1) into wild-type Col-0 using the floral dip method (Clough and Bent, 1998) and selected on medium containing 25 μg/mL hygromycin B.

### Confocal imaging for BiFC and subcellular localization analysis

Subcellular localization of Arabidopsis transgenic OMPL plants, BiFC analysis of protein pairs and subcellular localization of individual proteins or co-localization of protein pairs in *Nicotiana benthamiana* leaves were examined using confocal microscopy (Zeiss LSM 710). YFP and chlorophyll fluorescence were excited at 514 nm with detection ranges of 525–600 nm and 650–720 nm, respectively. MitoTracker Deep Red FM (Invitrogen M22426) was excited at 644 nm with a detection range of 578–696 nm. mCherry was excited at 561 nm, and its fluorescence was detected over the range 578–637 nm. CFP was excited at 405 nm, its fluorescence was detected over the range 454–487 nm.

### Growth assessment under stress

Arabidopsis seeds were surface-sterilized and sown on solidified half-strength MS medium. After stratification at 4°C for 2 d in the dark, the plates were placed vertically at 23°C, in a long-day (16-h light/8-h dark) photoperiod. Five-day-old seedlings were transferred to MS plates supplemented with or without the stressors 10 µM antimycin, 1 mM H_2_O_2_, 100 mM NaCl or 150 mM mannitol. These plates were placed at 22°C for 5–10 d under a long-day photoperiod as described above before the phenotypes were recorded by taking photographs and measuring primary root length and fresh weight.

### Optimization of biotin assays in Arabidopsis

For biotin treatment, whole Arabidopsis seedlings were grown vertically on half-strength MS medium containing 0.5% (w/v) sucrose for 14 days under long-day conditions (16-h light/8-h dark) at 22°C. About 0.1 g seedlings was pooled and incubated with biotin solution (0.5 μM, 50 μM or 100 μM biotin resuspended in water) in a nine-well plate with shaking (80–100 rpm) at room temperature (22°C) for 1 h or incubated with 100 μM biotin solution for 1, 2 or 3 h. Following treatment, the seedlings were dried with paper towels and flash-frozen for later immunoblotting. Comparison of the biotin ligase activity with different biotin concentrations or labeling duration was performed twice.

### Biotin labeling of control seedlings and seedlings exposed to UV-B or HL stress

CP-OMPL seedlings (14-day-old) grown vertically on half-strength MS plates were placed in a growth chamber under a light intensity of 2,000 μmol m^−2^ s^−1^ for 1 h. Fourteen-day-old MT-OMPL and PX-OMPL seedlings grown vertically on half-strength MS plates were treated with UV-B light (6 W/m^2^) for 6 h. All stress-treated seedlings were allowed to recover for 24 h under normal light and at room temperature before being incubated with biotin. About 2 g wild-type, stress-treated and untreated OMPL seedlings were incubated in 150-mL flasks containing 80 mL 100 μM biotin with shaking (80–100 rpm) at room temperature for 3 h. After biotin treatment, seedlings were quickly rinsed with ice-cold water and washed three times to stop the labeling reaction (Branon et al., 2018) and to remove excess biotin. The seedlings were then dried with paper towels, a sample of approximately 0.1 g seedlings was taken for immunoblots, and the remaining seedlings were pooled, frozen in liquid nitrogen, and stored at –80°C until further use. For affinity purification with streptavidin beads, one aliquot of plant material powder was resuspended in 3 mL ice-cold extraction buffer (50 mM Tris-HCl pH 7.5, 150 mM NaCl, 0.1% [w/v) SDS, 0.1% [v/v) Triton X-100, 0.1% [w/v) sodium deoxycholate, 1 mM EDTA, 10 mM NaF, 1 mM PMSF and 1x complete proteasome inhibitor), mixed well and incubated on a rotor wheel at 4°C for 10 min. The suspension was centrifuged for 10 min at 4°C and 15,000 g to remove cell debris before the supernatant was applied to a PD-10 desalting column (Thermo Fisher Scientific) to remove excess free biotin using the centrifugation protocol according to the manufacturer’s instructions. The column was equilibrated with five volumes of ice-cold equilibration buffer (extraction buffer without complete and PMSF), 2.5 mL protein extract was loaded and proteins were eluted with 3.5 mL equilibration buffer. The protein concentration of the protein extract was then measured with a BCA Protein Assay Kit (Thermo Fisher Scientific). A volume of each protein extract corresponding to 14 mg protein was transferred into a new 5-mL tube containing 150 µL streptavidin beads (Thermo Fisher Scientific) that were pre-washed with extraction buffer and incubated on a rotor wheel at 4°C overnight. The beads were washed with wash buffer (50 mM Tris-HCl pH 7.5, 500 mM NaCl, 1 mM EDTA, 10 mM NaF) three times. About 10 µL beads was boiled in 30 µL 4x Laemmli buffer at 95°C for 5 min for immunoblots to examine the success of the procedure.

### Sample preparation for MS analysis

For on-beads tryptic digest, frozen streptavidin beads were thawed and suspended in trypsin buffer (12 mM sodium deoxycholate and 12 mM sodium lauroyl sarcosinate in 100 mM Tris-HCl pH 8.5) and then were diluted 4-fold with 50 mM triethylammonium bicarbonate buffer (Thermo Fisher Scientific). Protein was then digested with 2.5 μg Lys-C (Wako, Japan) for 4 h at 37°C and 2 μg trypsin (Promega, V5113) for an additional 4 h at 37°C. After digestion, the digestion buffer and beads were separated by centrifugation at 12,000 g for 5 min at room temperature . The detergents in the supernatants were removed by acidifying the solution using 10% (w/v) trichlorofluoric acid and then centrifuged at 16,000 g for 20 min at room temperature. After desalting using a C18 StageTip, the supernatants were subjected to LC-MS/MS analysis.

### LC-MS/MS

Peptides were dissolved in 4 μL 0.2% (v/v) formic acid (FA) and injected into an Easy-nLC 1200 (Thermo Fisher Scientific). Peptides were separated on a 15-cm in-house packed column (360 μm OD × 75 μm ID) containing C18 resin (2.2 μm, 100 Å, Michrom Bioresources). The mobile phase buffer consisted of 0.1% (v/v) FA in ultra-pure water (Buffer A) with an eluting buffer of 0.1% (v/v) FA in 80% (v/v) acetonitrile (Buffer B) run over a linear 93-min gradient of 5–40% (v/v) buffer B at a flow rate of 300 nL/min. An Easy-nLC 1200 was coupled online with an Q Exactive HF-X (Thermo Fisher Scientific). The mass spectrometer was operated in data-dependent mode in which a full-scan MS (from m/z 350–2,000 with a resolution of 120,000) was followed by top 10 higher-energy collision dissociation MS/MS scans of the most abundant ions with dynamic exclusion for 45 s.

### Proteomics data analysis

Proteins were identified and quantified in Proteome Discoverer software (version 2.4) using SEQUEST HT search algorithm. Peptides were searched against the latest TAIR10 protein database containing 35,386 entries (TAIR10_- pep_20101214, updated 2011-08-23, www.arabidopsis.org). Peptide precursor mass tolerance was set to 10 ppm, and MS/MS tolerance was set to 0.02 Da. Three missed cleavage sites of trypsin were allowed. The false discovery rate of proteins and peptides was set to 1%. Unique and razor peptides were used for protein quantification.

### Identification of enriched proteins in the OM proximity proteome

To remove nonspecific labeling and define the OM proximity proteome, we applied a filtering to the pairwise comparisons of NMT vs WT, NPX vs WT, and NCP vs WT samples using only proteins that were found in three replicates of the corresponding NMT, NPX, NCP and WT samples and criteria of FC > 1.5 and *p* < 0.05. For UV-B- and HL stress-treated samples, we applied a filtering to the pairwise comparisons of UMT vs NMT, UPX vs NPX, and HL24 vs NCP samples for the proteins that were only found in two replicates of NCP and HL24 samples (Fig. 4A) or two and three replicates of NMT and UMT or NPX and UPX samples (Fig. 4B) and criteria of |FC| > 1.5 and *p* < 0.05.

### Y2H and Co-IP assays

The yeast strain Y2H was co-transformed with the relevant constructs and grown on synthetic defined (SD) medium lacking Trp and Leu (SD –Trp –Leu) for selection of transformants. Positive colonies were incubated in SD –Trp –Leu liquid medium at 28°C, and serial dilutions were spotted onto SD –Trp –Leu –His medium plates. For Co-IP, about 1 g *N. benthamiana* leaves that were co-infiltrated with the appropriate pairs of constructs was ground in liquid nitrogen, and the resulting powder was resuspended in 0.5 mL extraction buffer (50 mM HEPES, 100 mM NaCl, 10 mM EDTA, 10% [v/v] glycerol, 0.2% [v/v] NP-40, 0.5 mM DTT, 1 mM PMSF, and 1 protease inhibitor cocktail tablet/50 mL, pH 7.5) and centrifuged at 12,000 g for 20 min at 4°C. Anti-cMyc (Chromotek, 004282-07-02) or HA beads (MCE, HY-K0201) were added to the protein extracts and incubated at 4°C for 4 h. Beads-bound proteins were collected by centrifugation at 12,000 g for 1 min at 4°C and washed four times with extraction buffer, boiled in 1× SDS sample buffer, and run on an SDS-PAGE gel for immunoblotting with antibodies (Dataset S5).

### GO enrichment analysis

The PANTHER website was used for GO term enrichment analysis of OM proximity proteomes (|FC| > 1.5, *p* < 0.05). ImageGP was used to generate GO enrichment plots, PCA analysis and cluster correlation plots and heatmaps.

## Data availability

The proteomics data were deposited to the ProteomeXchange (PXD039752) and JPST002016 for jPOST (URL:https://repository.jpostdb.org/preview/129213289763da3a6be1611, Access key: 5013).

## AUTHOR CONTRIBUTIONS

X.Z. conceived the project and wrote the manuscript. X.B., H.J. and X.Z. generated constructs and performed PL experiments and data analysis as well as confocal imaging analysis. Y.Z. conducted the Y2H assay. X.L., P.L. and C.M. assisted with experiments. X.Z., P.W. and C.-P.S. analyzed the data.

## Supplemental Materials

**Supplemental Dataset S1.** OM proximity proteomes and GO analysis.

**Supplemental Dataset S2.** Mitochondrial OM proximity proteome after UV-B stress and GO analysis.

**Supplemental Dataset S3.** Peroxisome OM proximity proteome after UV-B stress and GO analysis.

**Supplemental Dataset S4.** Chloroplast OM proximity proteome after HL stress and GO analysis.

**Supplemental Dataset S5.** Primer sets and antibodies.

**Supplemental Table S1**. A list of OMPL constructs generated in this study.

**Supplemental Table S2.** A list of OM receptor interactors detected by BiFC assay.

**Supplemental Figure S1.** Design of the expression cassettes for targeting biotin ligase to cytosolic-facing mitochondrion, peroxisome and chloroplast outer membrane.

**Supplemental Figure S2.** Transgenic Arabidopsis seedlings harboring OMPL constructs show normal growth under normal and stress conditions.

**Supplemental Figure S3.** Subcellular localization of TOM20-2 and NAC4.

**Supplemental Figure S4.** UV-B and HL stress alters the OM proximity proteome.

**Supplemental Figure S5.** NAC4 shows no or weak interaction with ribosomal proteins and TOM20-2 proteins.

**Supplemental Figure S1. Design of the expression cassettes for targeting biotin ligase to cytosolic-facing mitochondrion, peroxisome and chloroplast outer membrane.** (A-D) Schematic diagrams of the expression cassette encoding the PL enzymes biotin ligase TurboID (A and B) or minTurboID (C and D). MT, PX and CP are mitochondrion, peroxisome and chloroplast outer membrane anchored proteins, respectively; EGFP, enhanced green fluorescence protein between the PL enzyme and outer membrane anchored protein to add distance between the PL enzyme and the outer membrane (TR and TRm) (B and D), or to the N-terminal or C-terminal end of the PL enzymes to allow a direct link between the PL enzyme and the outer membrane (T and Tm) (A and C) for visualizing the subcellular localization of PL enzymes. HA, hemagglutinin tag.

**Supplemental Figure S2. Transgenic Arabidopsis seedlings harboring OMPL constructs show normal growth under normal and stress conditions.** (A, C, E, G) Overall morphology of OMPL seedlings. Five-day-old seedlings transferred to MS medium alone or containing stressors as indicated. Photographs were taken 7 days after transfer. Scale bar, 1 cm. (B, D, F, H) Root length and fresh weight of OMPL seedlings. Root length and fresh weight of three biological replicates were measured and no significant differences were detected. MT-OMPL seedlings are shown in A and B; PX-OMPL seedlings in C-F; CP-OMPL seedlings in G and H. The lines used in this study are labeled in red.

**Supplemental Figure S3. Subcellular localization of TOM20-2 and NAC4.** (A) *YFP-TOM20*-2 or *YFP-NAC4* were transiently co-expressed with a peroxisome marker construct in *N. benthamiana* leaves. (B) *YFP-TOM20*-2 or *YFP-NAC4* were transiently co-expressed with a mitochondrial marker construct in *N. benthamiana* leaves. Arrows indicate YFP merged with mitochondrial or peroxisome marker signals. Scale bars, 20 μm.

**Supplemental Figure S4. UV-B and HL stress alters the OM proximity proteome.** (A) Venn diagram analysis of OMPL proteins becoming more abundant or less abundant upon UV-B, or HL stress in UMT, UPX and HL24 samples. (B, C) Heatmap representation of the abundance of ribosomal proteins with significant reduction (B) or increase (C) in abundance in UMT samples. (D) Heatmap representation of the abundance of ribosomal proteins with significant reduction in abundance in UPX samples. (E) GO term analysis of stress-increased OMPL proteins.

**Supplemental Figure S5. NAC4 shows no or weak interaction with ribosomal proteins and TOM20-2 proteins.** BiFC assay showing that NAC4 weakly interacts with RPL11 and TOM20-2 in the cytosol, but not with RPL5 and RPL13. Scale bar, 20 μm.

